# Ethacridine inhibits SARS-CoV-2 by inactivating viral particles in cellular models

**DOI:** 10.1101/2020.10.28.359042

**Authors:** Xiaoquan Li, Peter Lidsky, Yinghong Xiao, Chien-Ting Wu, Miguel Garcia-Knight, Junjiao Yang, Tsuguhisa Nakayama, Jayakar V. Nayak, Peter K. Jackson, Raul Andino, Xiaokun Shu

**Affiliations:** Department of Pharmaceutical Chemistry, University of California, San Francisco, San Francisco, California, USA; Cardiovascular Research Institute, University of California, San Francisco, San Francisco, California, USA; Department of Microbiology and Immunology, University of California, San Francisco, San Francisco, California, USA; Department of Baxter Laboratory for Stem Cell Biology, Department of Microbiology & Immunology, Stanford University, California, USA; Department of Otolaryngology–Head and Neck Surgery, Stanford University, Stanford, California, USA

## Abstract

SARS-CoV-2 is the coronavirus that causes the respiratory disease COVID-19, which is now the third-leading cause of death in the United States. The FDA has recently approved remdesivir, an inhibitor of SARS-CoV-2 replication, to treat COVID-19, though recent data from the WHO shows little to no benefit with use of this anti-viral agent. Here we report the discovery of ethacridine, a safe antiseptic use in humans, as a potent drug for use against SARS-CoV-2 (EC_50_ ~ 0.08 *μ*M). Ethacridine was identified via high-throughput screening of an FDA-approved drug library in living cells using a fluorescent assay. Interestingly, the main mode of action of ethacridine is through inactivation of viral particles, preventing their binding to the host cells. Indeed, ethacridine is effective in various cell types, including primary human nasal epithelial cells. Taken together, these data identify a promising, potent, and new use of the old drug possessing a distinct mode of action for inhibiting SARS-CoV-2.

## Introduction

The worldwide outbreak of the respiratory disease COVID-19 is caused by the coronavirus SARS-CoV-2 (severe acute respiratory syndrome coronavirus 2). SARS-CoV-2 is an RNA betacoronavirus of the family *Coronaviridae*. It contains a single-stranded positive-sense RNA genome encapsulated within a membrane envelope ^1-4^. Its genomic RNA is approximately 30kb with a 5′-cap structure and 3′-poly-A tail ^1^. The genome of SARS-CoV-2 can be split into two main regions that contain as many as 14 open reading frames (ORFs) ^5^.

The first region, containing the first ORF (ORF1a/b), is about two-thirds of the length of the genome. After coronavirus attachment and entry into the host cell, the viral genomic RNA is released intracellularly. The first region of the genomic RNA is used as a template to directly translate two polyproteins: pp1a and pp1ab. The pp1a polyprotein is translated from ORF1a. The pp1ab polyprotein comes from a -1 ribosomal frameshift between ORF1a and ORF1b. The two overlapping polyproteins are processed by a papain-like protease (PLpro) and a 3-chymotrypsin-like protease (3CLpro). Both pp1a and pp1ab are mainly processed by 3CLpro, which is also referred to as the main protease (Mpro). Mpro digests the polyproteins in at least 11 conserved sites, starting with the auto-proteolytic cleavage of this viral protease itself from pp1a and pp1ab. The functional polypeptides, 16 non-structural proteins (nsp1–16), are released from the polyproteins after extensive proteolytic processing. Nsp12 (i.e., RNA-dependent RNA polymerase (RdRp)), together with other nsps (e.g., nsp7 and nsp8), forms a multi-subunit replicase/transcriptase complex (RTC) that is associated with the formation of virus-induced double-membrane vesicles ^4,6,7^. The membrane-bound RTC synthesizes a full-length negative-strand RNA template that is used to make positive-strand viral genomic RNA.

The remaining one-third of the genomic RNA is used by the RTC to synthesize subgenomic RNAs (sgRNAs) that encode four conserved structural proteins (spike protein (S), envelope protein (E), membrane protein (M), and nucleocapsid protein (N)), and several accessory proteins. Eventually, the viral RNA-N complex and S, M, and E proteins are assembled in the ER-Golgi intermediate compartment (ERGIC) to form a mature virion that is then released via budding from the host cell. S protein is exposed on the surface of the virion and binds the virus receptor ACE2 on the host cell surface. Therefore, Mpro plays a central and critical role in the lifecycle of the coronavirus and is an attractive drug target ^8-10^, which also include other biological steps essential for viral replication and budding.

To identify drugs that may inhibit the coronavirus, we redesigned the green fluorescent protein (GFP) into an activity reporter of Mpro, which becomes fluorescent only upon cleavage by the active Mpro. Using this fluorescent assay, we screened a drug library in living cells and identified several drugs that inhibit Mpro activity. One highly effective drug, ethacridine, inhibits SARS-CoV-2 production by inactivating the viral particles.

## RESULTS

### Rational design of a fluorogenic Mpro activity reporter FlipGFP^Mpro^

To develop an activity reporter of Mpro with a large dynamic range suitable for high-throughput screening (HTS), we applied the GFP-based protease reporter called FlipGFP ^11^, which was designed by flipping one of the 11 beta-strands of a split GFP. Briefly, the split GFP contains two parts: one part contains beta-strands 10 and 11 (i.e., GFP10 and 11), and the other contains nine other beta-strands and the central alpha helix (i.e., GFP1– 9). GFP10–11 contains the highly conserved Glu222 that is essential for catalyzing chromophore maturation. GFP1-9 contains the three amino acids that form the chromophore via cyclization, dehydration and oxidation ^12^. GFP10-11 spontaneously binds GFP1-9 and catalyzes the chromophore maturation, leading to green fluorescence. To design an Mpro activity reporter, we “flipped” GFP10-11 using heterodimeric coiled coils (E5 and K5) so that the flipped GFP10-11 cannot bind GFP1-9 when Mpro is inactive, and thus, no or little fluorescence is detected (**Fig. 1a**). We incorporated an Mpro-specific cleavage sequence AVLQ↓SGFR (↓ denotes the cleavage site) between GFP11 and K5. In this way, when Mpro cleaves the proteolytic site, GFP11 is flipped back, allowing GFP10-11 to now bind GFP1-9, resulting in bright fluorescence (**Fig. 1a**). We named this reporter FlipGFP^Mpro^. To normalize the fluorescence, we added a red fluorescent protein mCherry within the construct via a “self-cleaving” 2A peptide^13^ (**Fig. 1c**).

**Fig. 1.**
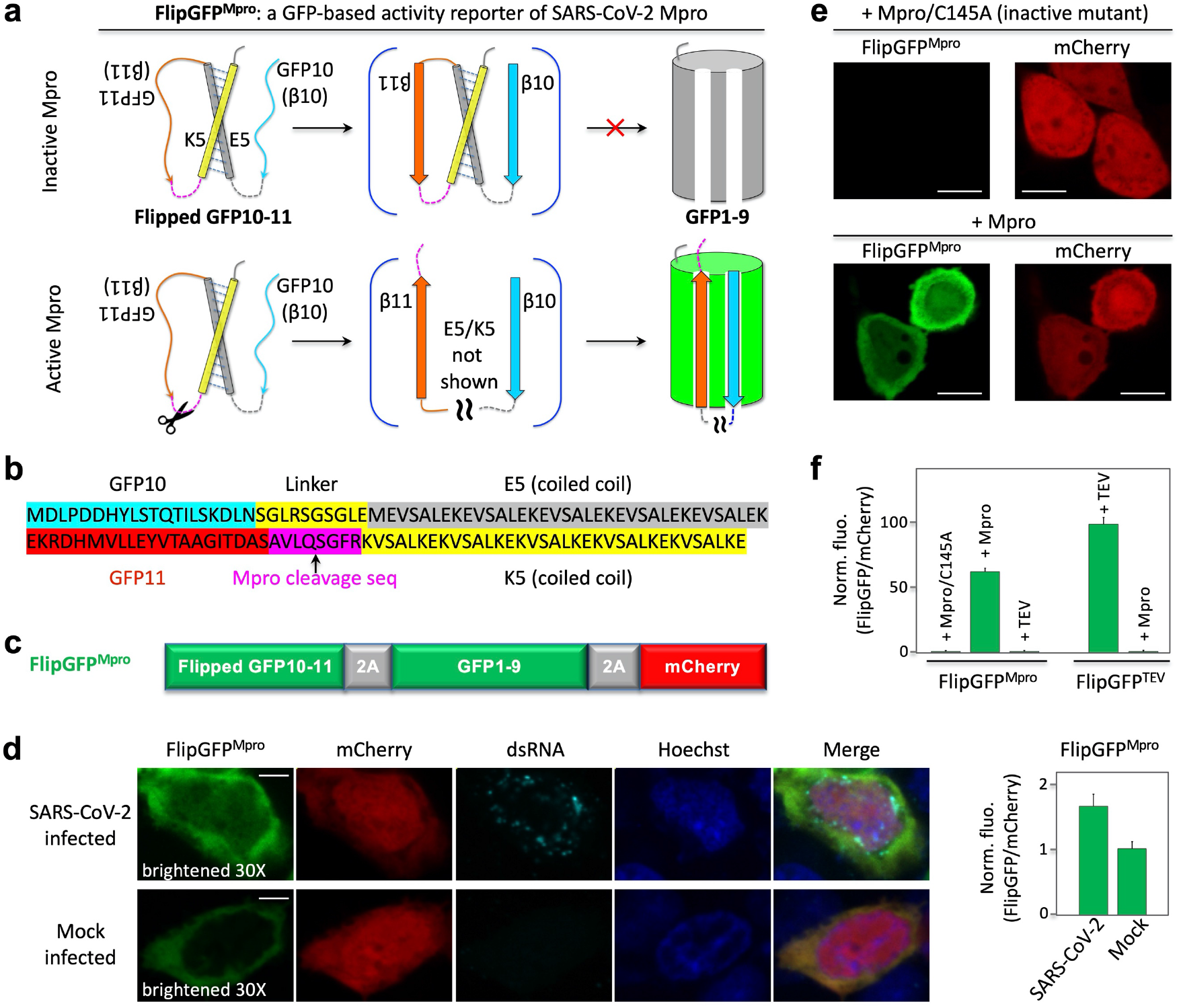
Design and demonstration of a GFP-based activity reporter of SARS-CoV-2 main protease Mpro. (**a**) Schematic of the reporter. (**b**) Sequence of the flipped GFP10-11. (**c**) Construct of the reporter FlipGFP^Mpro^. (**d**) Fluorescence images (left) and quantitative analysis (right) of SARS-CoV-2 or mock-infected HEK293T cells that co-expressed hACE2. The images in the FlipGFP channel were brightened 30-fold compared to those in (**e**). (**e**) Fluorescence images of HEK293T cells expressing FlipGFP^Mpro^ and mCherry, together with the inactive Mpro mutant C145A (upper panels) or wild type Mpro (lower panels). (**f**) Normalized FlipGFP fluorescence by mCherry. The ratio of FlipGFP/mCherry for the Mpro/C145A is normalized to 1. Data are mean ± SD (n = 5). FlipGFP^TEV^ is a TEV activity reporter containing TEV cleavage sequence in FlipGFP. Scale bar: 5 *μ*m (**d**); 10 *μ*m (**f**).

To determine if FlipGFP^Mpro^ serves to report on Mpro activity of SARS-CoV-2 in living cells, we expressed the human angiotensin converting enzyme 3 (ACE2), the SARS-CoV2 receptor, in HEK293-FlipGFP^Mpro^ cells. Next, we infected the cells with SARS-CoV-2, and at 24 hours post-infection, cells were analyzed by immunofluorescence using antibodies directed against double-stranded RNA (dsRNA) and FlipGFP^Mpro^ green fluorescence. The green fluorescence of the sensor, normalized by the co-expressed mCherry, was 63% greater in the coronavirus-infected cells than in mock-infected cells (Fig. 1d). Infected cells also showed dsRNA fluorescence compared to non-infected (mock) cells without dsRNA staining (Fig. 1d). These data demonstrate that the utility of FlipGFP^Mpro^ as a reporter of SARS-CoV-2 Mpro activity in human cells.

Next, we established a system for screening Mpro inhibitors in living cells by exogenously expressing Mpro in HEK293. Specifically, wild-type Mpro or an inactive Mpro mutant (with catalytic cysteine 145 mutated to alanine) were co-expressed in this cell line. The green fluorescence of FlipGFP^Mpro^ was barely detected in the cells expressing the inactive Mpro/C145A mutant, whereas the red fluorescence of mCherry was observed (**Fig. 1e**, upper panels). On the other hand, strong green fluorescence was detected in the cells expressing Mpro with similar levels of mCherry fluorescence (**Fig. 1e**, lower panels). The green fluorescence of FlipGFP^Mpro^, normalized to the red fluorescence of mCherry, revealed an ∼60-fold dynamic range between inactive and active Mpro (**Fig. 1f**). Furthermore, based on these quantified data, we calculated a *Z*’*-factor*^14^

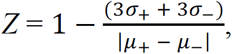

which is ∼0.8 with Mpro and its inactive mutant as positive (+) and negative (-) controls, respectively (here σ is standard deviation, *μ* is mean). This suggests that the assay is robust for HTS.

The FlipGFP^Mpro^ sensor was not responsive to the TEV protease, and the FlipGFP-based TEV reporter (FlipGFP^TEV^) was only activated by TEV but not by Mpro (**Fig. 1f**). Thus, FlipGFP^Mpro^ specifically detects Mpro activity with a large dynamic range. Therefore, our data show that we have established a robust HTS system for screening Mpro inhibitors at a BSL2 level with 60-fold fluorescence change and a robust z’-factor. Comparing the normalized FlipGFP^Mpro^ fluorescence in the SARS-CoV-2 infected cells (**Fig. 1d**) with that from the cells expressing Mpro exogenously (**Fig. 1e, f**) suggests that the active Mpro concentration in the cytoplasm of the coronavirus-infected cells is ∼100-fold lower than that of the HEK293 cells exogenously expressing Mpro (under a EF1*α* promoter).

### HTS of drugs that inhibit Mpro activity in living cells

Next, we conducted HTS of ∼1600 FDA-approved drugs (20 *μ*M final concentration, **Fig. 2a**). The reporter construct (FlipGFP^Mpro^ and mCherry) was transfected into HEK293 cells, followed by addition and incubation of the drugs. Levels of green fluorescence normalized to red fluorescence were calculated. A volcano plot revealed ∼120 drugs that showed ≥ 50% reduction of Mpro activity with a p-value < 0.001 (**Fig. 2a**). To confirm this result, we re-screened the identified ∼120 drugs under similar conditions (Supporting Fig. S1). We further assayed those top 15 drugs at a lower concentration (10 *μ*M) and found that 12 drugs showed ≥50% reduction of FlipGFP^Mpro^ fluorescence (normalized by mCherry) at 10 *μ*M concentration (**Fig. 2b**). We finally calculated an IC_50_ for each of the 12 drugs. Six drugs were at 2 – 6 *μ*M (**Fig. 2c**), and the rest were above 6 *μ*M (Supporting Fig. S2).

**Fig. 2.**
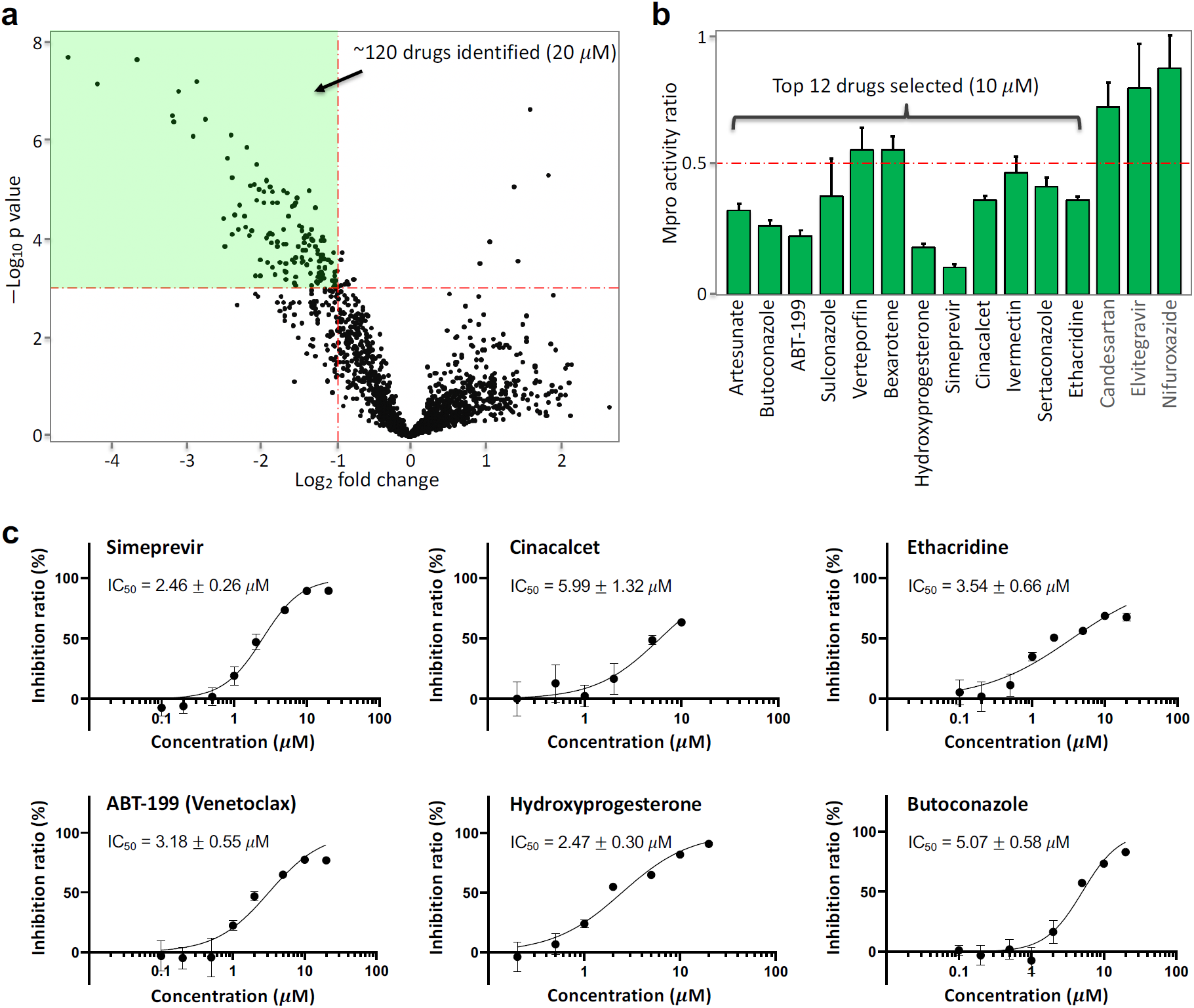
High-throughput screening and drug identification using FlipGFP^Mpro^ in living cells. (**a**) Volcano plot of 1622 FDA-approved drugs (20 *μ*M) in inhibiting Mpro. HEK293T cells were transfected with FlipGFP^Mpro^, mCherry and Mpro. FlipGFP fluorescence was normalized to co-expressed mCherry. (**b**) Normalized ratio of Mpro activity in drug (10 *μ*M) vs DMSO-incubated cells for the third-round validation. The Mpro activity was determined as FlipGFP fluorescence normalized to mCherry. The ratio of Mpro activity was calculated by normalizing Mpro activity with that of cells treated with DMSO. Data are mean ± SD (n = 5). The 15 drugs were identified from a second-round imaging of the 120 identified drugs (20 *μ*M, Extended data Fig. 1). (c) Dose-response curve of top six drugs in inhibiting Mpro. Inhibition ratio was calculated as (1-(ratio of Mpro activity)) × 100%. IC_50_ was represented as mean ± SEM (n = 5). See Extended data Fig. 2 for the other six drugs.

### Antiviral activity of identified drugs

We next investigated antiviral activity of selected drugs in Vero E6 cells. The cell monolayers were pretreated with the 12 selected drugs for 3 hours, and then infected with SARS-CoV-2. The cells were further cultivated in the presence of each respective compound at a concentration of 5 *μ*M. After 16 hours of incubation, the culture media samples were collected, and the amount of infectious particles were estimated by plaque assay (**Fig. 3a, b** & Supporting Fig. S3). Our data revealed that 9 of the 12 drugs showed significant antiviral activity at 5 *μ*M. Strong inhibition was detected for ethacridine with 5 – 6 logs reduction in viral titer, simeprevir ∼4-log reduction, ABT-199 ∼2-log reduction, hydroxyprogesterone ∼1-log reduction, cinacalcet ∼1-log reduction. Two of the 12 drugs (ivermectin and verteporfin) were cytotoxic at 5–13 *μ*M in Vero E6 cells and, thus, were excluded from the further analysis (Supporting Fig. S4). As a comparison, we tested the antiviral effect of a reported Mpro inhibitor, ebselen, which showed ∼2-log reduction in viral titer. The RdRp inhibitor remdesivir showed ∼4-log reduction.

**Fig. 3.**
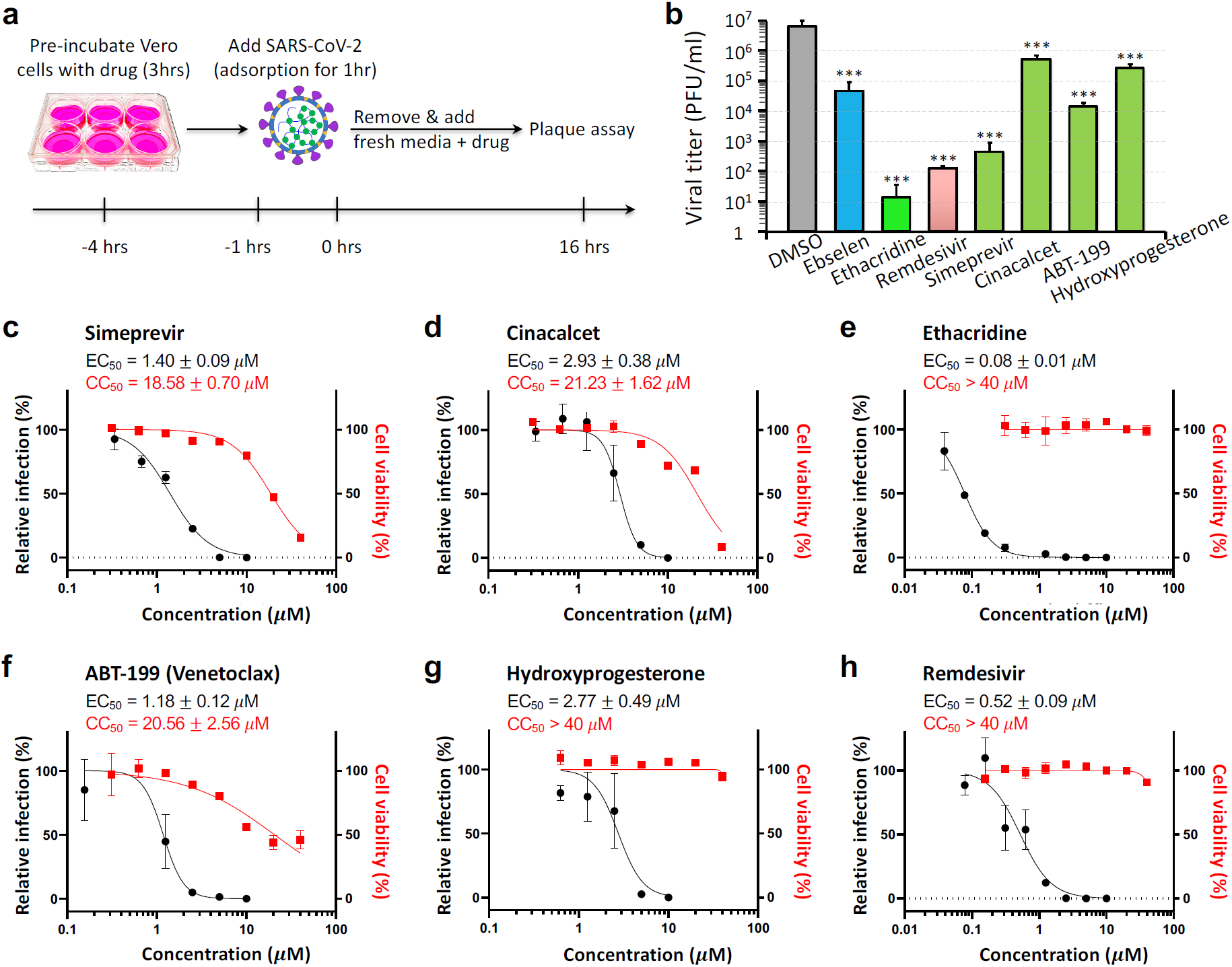
Antiviral activities of the identified drugs against SARS-CoV-2. (**a**) Schematic showing the experimental design, see methods for details. (**b**) Quantitative analysis of viral titer from plaque assays on Vero E6 cells treated with each drug at 5 *μ*M. Data are mean ± SEM (n = 3). *** *p* < 0.001. (**c–h**) Dose-response and cell-toxicity curve of each drug against SARS-CoV-2 by plaque assays. The percentage of relative infection was determined by the ratio of infection rate of SARS-CoV-2 treated with each drug divided by that of DMSO control. EC_50_ and CC_50_ are represented as mean ± SEM (n = 3).

Next, we determined dose-response curves for the top 5 selected drugs. First, our data revealed that the EC_50_ of four drugs (simeprevir, cinacalcet, ABT-199 and hydroxyprogesterone) was 1–3 *μ*M (**Fig. 3c – g**), within a range similar to their IC_50_ in inhibiting Mpro (**Fig. 2c**). This was consistent with the expectation that SARS-CoV-2 replication is inhibited by restricting Mpro activity. Indeed, as we were finalizing this study, a preprint report showed that simeprevir inhibits Mpro activity and SARS-CoV-2 ^15^. By contrast, ethacridine showed outstanding antiviral activity (EC_50_ ∼ 0.08 *μ*M, **Fig. 3e**), which is 40-fold lower (i.e. stronger) than its Mpro-inhibiting activity (IC_50_ ∼ 3.54 *μ*M, **Fig. 2c**). These data suggest that the outstanding antiviral activity of ethacridine is not mainly accounted for by its Mpro-inhibiting activity. Lastly, for comparison, we also determined the EC_50_ of remdesivir ∼0.52 *μ*M in a side-by-side manner (**Fig. 3h**), which indicates that has more potent antiviral activity than remdesivir.

### Ethacridine inhibits SARS-CoV-2 by inactivating viral particles

To determine how ethacridine inhibits SARS-CoV-2, we tested infectivity of the virus particles after ethacridine treatment using plaque assay, and we also measured viral RNA levels using qRT-PCR. We examined the antiviral effect of ethacridine on different stages of the viral lifecycle of SARS-CoV-2, including before virus-cell binding, after cell entry, and after budding. In particular, we pre-incubated SARS-CoV-2 particles with ethacridine (5 µM) or DMSO for 1 hour. The mixture was then added to Vero E6 cells for viral adsorption at a multiplicity of infection (MOI) at 0.5. Next, we removed the solution and added fresh medium containing ethacridine (5 µM) or DMSO. Sixteen hours later, we collected supernatant and conducted plaque assay with overlaid agar without ethacridine or DMSO, measured viral titer of the supernatant. We also conducted qRT-PCR and measured viral RNA levels in the supernatant and within cells. In this way, we developed three conditions (**Fig. 4a**): 1) Control (DMSO + DMSO): the virus and cells were not exposed to DMSO and not the drug; 2) The virus and cells were exposed to the drug at all stages, including 1 hour before infection, during replication, and after viral budding (i.e. Eth. + Eth.); 3) The virus and cells were exposed to the drug only after viral entry, during replication, and after budding (i.e. DMSO + Eth.). Lastly, we used a fourth condition (**Fig. 4b**): we conducted plaque assay right after pre-treatment of SARS-CoV-2 with ethacridine for 1 hr (i.e. Eth. [1 hr]), which determines antiviral activity of the drug on the viral particles (equivalent to addition of the drug after viral budding).

**Fig. 4.**
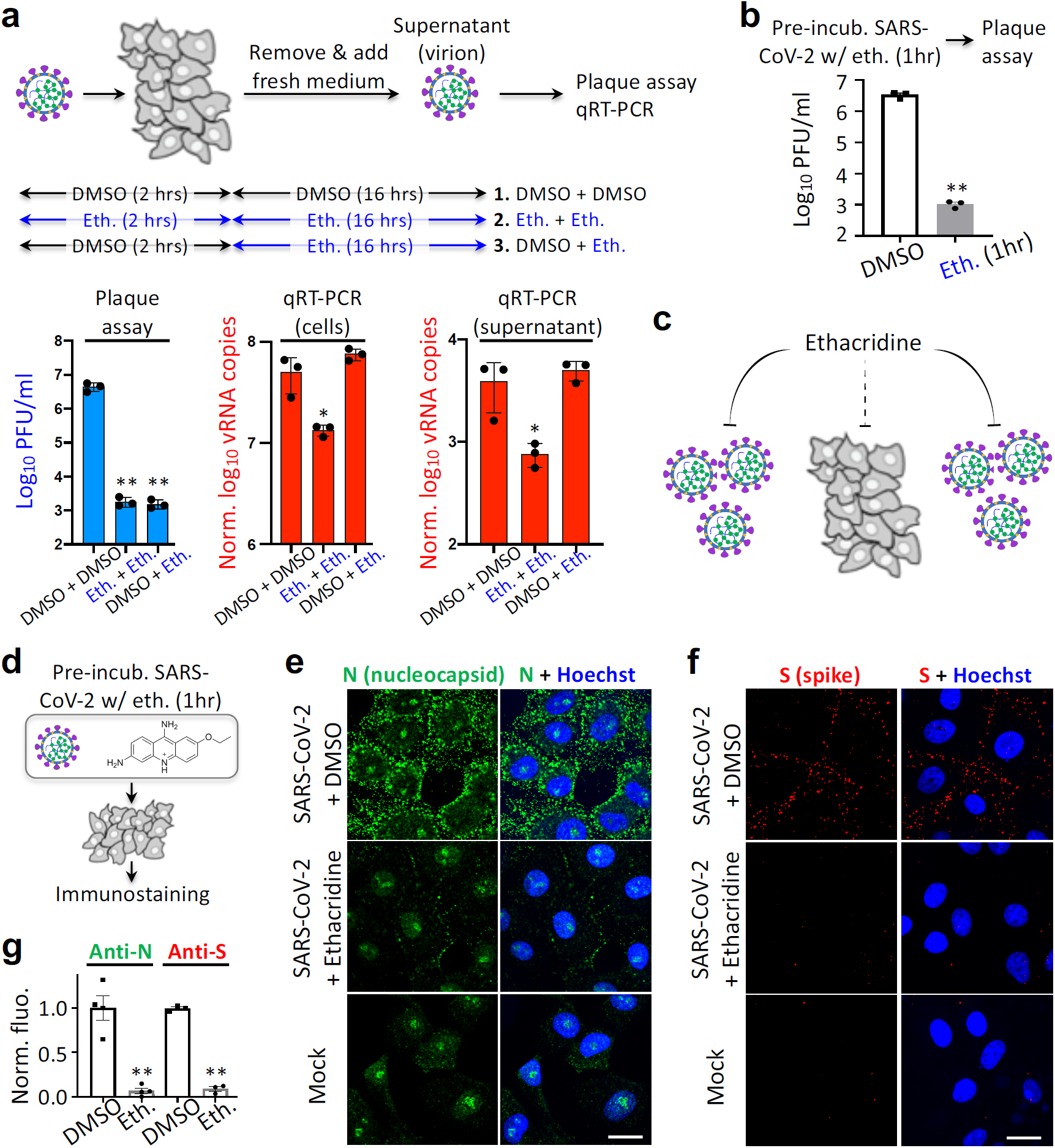
Ethacridine bocks SARS-CoV-2 by inactivating viral particles. (**a**) Upper panel: schematic showing the experimental design for plaque assay and qRT-PCR. The virus was pre-incubated with ethacridine or DMSO for 1 hr. The mixture was added to Vero cells for adsorption at 4°C for 1 hr. Details can be found in the text. Lower panel: quantitative analysis of viral titer from plaque assay (left), and viral RNA (vRNA) copies by qRT-PCR in Vero cells (middle) and supernatant (right). (**b**) Quantitative analysis of viral titer by plaque assay. (**c**) Proposed mode of action of ethacridine by mainly inactivating viral particles of the coronavirus with no or little effect on viral RNA replication. (**d**) Schematic showing the experimental design for immunostaining. (**e, f**) Representative images of immunostaining against nucleocapsid protein (N) and spike (S) in Vero E6 cells after infection with the virus that was pre-treated with ethacridine (5 *μ*M) or DMSO (control), or no infection (mock). (**g**) Quantitative analysis of the immunofluorescence. Data are mean ± SEM (n = 3 or 4 biological replicas). **: p value < 0.01. *: p value < 0.05. Scale bar, 20 *μ*m.

When ethacridine was present continuous prior to plaque assay (Eth. + Eth.), viral titers were reduced 3 – 4 logs, compared with the control (DMSO + DMSO) (**Fig. 4a**, lower left panel). When ethacridine was added to the Vero cells after viral entry (DMSO + Eth.), similar level of reduced infectivity (3 – 4 logs reduction in viral titer) was observed (**Fig. 4a**). Interestingly, when SARS-CoV2 was pre-incubated with ethacridine for 1 hr (Eth. [1 hr]) and followed by plaque assay without drug, we also observed 3 – 4 logs reduction in infectivity (**Fig. 4b**). This result suggests that the drug directly inactivates SARS-CoV2 viral particles. Because of similar-level reduction in infectivity in all of the three conditions, our data strongly suggests that ethacridine inhibits SARS-CoV-2 mainly by inactivation of viral particles.

We next examined viral RNA accumulation in infected cells. qRT-PCR measurement revealed no change of viral RNA (vRNA) levels when the drug was added after viral binding and cell entry (DMSO + Eth.) in both the supernatant and the cells (**Fig. 4a**, lower middle and right panels), compared with the control (DMSO + DMSO). This indicates that the drug has no effect on vRNA replication. Because plaque assay-based measurement of the same-conditioned sample (DMSO + Eth.) showed 3 – 4 logs reduction in infectivity, our data suggests that ethacridine inhibits SARS-CoV-2 by inactivating the viral particles without effect on vRNA replication. This is consistent with the results of plaque assays for the supernatant samples with 3 different conditions that showed similar level of reduction in infectivity. Here the virions in the supernatant were exposed to the drug before plaque assay-based measurement of viral titer.

Next, when ethacridine was present continuously (i.e. Eth. + Eth.), 4 – 5 fold reductions were observed in vRNA copies in the supernatant and wthin the cells (**Fig. 4a**, middle and right panels). Because plaque assay-based measurement of the same conditioned sample (Eth. + Eth.) showed 2400-fold reduction in viral titer, the effect of ethacridine on viral replication (4 – 5 fold reduction) is about 500-fold smaller than its effect on viral infectivity. This further supports the conclusion that ethacridine inhibits SARS-CoV-2 by inactivating the viral particles. The 4 – 5 fold reduction of vRNA copies is likely due to reduced viral copy numbers that may bind to the cells (see below), because here the additional step is that the viruses were pre-incubated with the drug.

Thus, our data based on plaque assay and qRT-PCR of different conditioned samples suggests that ethacridine inhibits SARS-CoV-2 by mainly inactivating viral particles, including the virus before binding to Vero cells, as well as virions in the supernatant after budding from host cells, with no or little effect on vRNA replication (**Fig. 4c**).

To further investigate the mechanism of ethacridine-based inactivation of the viral particles, we conducted immunofluorescence staining and imaging to determine whether the ethacridine-treated SARS-CoV-2 can bind to and enter the cells. We treated SARS-CoV-2 with ethacridine (5 *μ*M) or DMSO for 1 hour. Then the virus was added to cells for adsorption (37°C, 1 hour) at a MOI = 100. Cells were then quickly washed and fixed with 4% PFA (**Fig. 4d**). Immunostaining with antibodies against the nucleocapsid protein (N) of SARS-CoV-2 showed strong anti-N fluorescence on the plasma membrane of the cells infected with virus+DMSO mixture, but little anti-N fluorescence in cells treated with virus+ethacridine mixture (**Fig. 4e**). Immunostaining against the Spike protein (S) of SARS-CoV-2 also revealed fluorescence signals on control cells, but minimal fluorescence on cells infected with the virus+ethacridine mixture (**Fig. 4f**). Quantification of fluorescence revealed that ethacridine treatment led to a dramatic reduction in detectable anti-N and anti-S fluorescence (**Fig. 4g**). These results indicate that ethacridine-treated SARS-CoV-2 cannot bind cells to initiate infection.

We also tested the dependency of the viral-inactivation effect of ethacridine on dose, incubation time and incubation temperature with a plaque assay. For the conditions tested, the viral-inactivation effect showed dose-dependency but was comparable to a 1- or 2-hour incubation at room temperature or 37°C (Supporting Fig. S5).

### Further validation in human cells

We further evaluated the anti-viral effect of ethacridine in human cells, including a human lung epithelial A549 cell line stably expressing human ACE2 (A549^ACE2^) and human primary nasal epithelial (HNE) cells to ensure that the antiviral effect is not restricted to the Vero E6 cells. A plaque assay in A549^ACE2^ cells revealed that ethacridine-treated virus showed a dramatic decrease in infectivity when applied to A549^ACE2^ cells (Supporting Fig. S6), indicating that the the effect of virus inactivation of ethacridine is not specific to Vero cells. This is consistent with the main mode of action of ethacridine that it blocks SARS-CoV-2 by inactivating the viral particles and thus its effect is independent of cell type. To further support this and validate the antiviral effects, primary HNE cells were infected with ethacridine-treated virus and incubated with the media containing 5 *μ*M ethacridine for 48 hours (**Fig. 5a**). The cilia organelles within the nasal epithelium have been shown to strongly and specifically express the ACE2 receptor exploited by SARS-CoV-2 ^16^. As a control, we used DMSO solution without ethacridine. Immunostaining using anti-N and anti-S antibodies in HNE cells showed that, in the presence of ethacridine, the proportion of infected primary nasal cells was dramatically decreased (**Fig. 5b, c**), suggesting that ethacridine protected HNE cells from viral infection. These data suggest that the antiviral effect of ethacridine is cell-independent, consistent with the main mode of action of this drug.

**Fig. 5.**
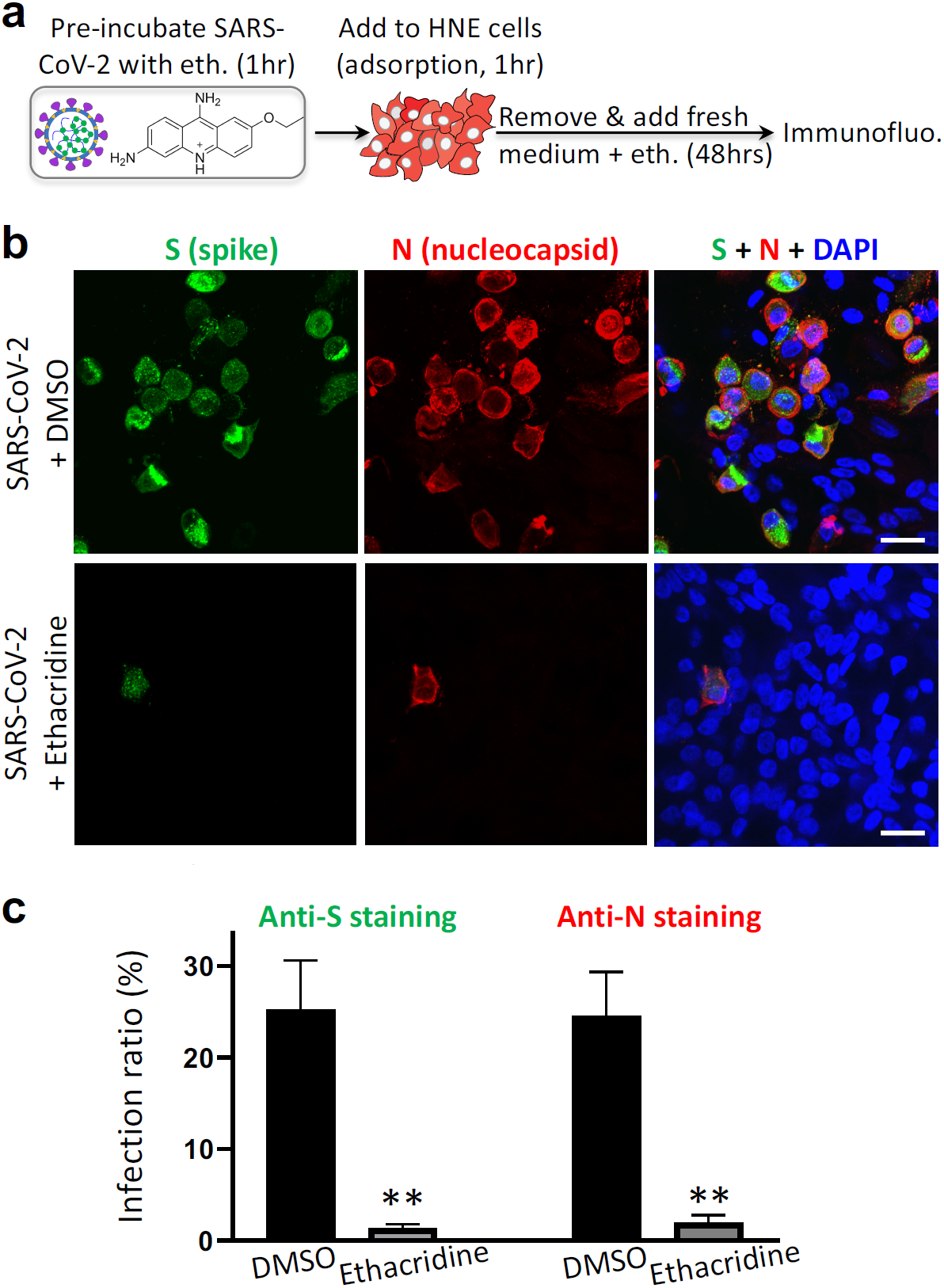
Ethacridine blocks SARS-CoV-2 in primary human nasal epithelial cells. (**a**) Schematic showing the experimental design, see methods for details. (**b**) Representative images of immunostaining against spike (S) and nucleocapsid protein (N) in HNE cells 48 hours after viral infection in the presence ethacridine (5 *μ*M) or DMSO (control), or no infection (mock). (**c**) Quantification of infection ratio based on anti-S or anti-N staining. Data are mean ± SEM (three biological replicates). **: p value < 0.01. Scale bar, 20 *μ*m.

## DISCUSSION

Several small molecule-based inhibitors have been identified that interfere with major targets involved in the viral lifecycle of SARS-CoV-2, including the main protease Mpro and the SARS-CoV-2 replicase RdRp. For example, *α*-ketoamide inhibitors of Mpro and the FDA-approved drug ebselen have been reported to inhibit Mpro activity with EC50 at 0.53 and 4.67 *μ*M, respectively ^8-10^. The replicase RdRp inhibitor, remdesivir, has been designed and very recently approved by FDA to treat critically-ill COVID-19 patients ^6,17^. However, very recent data from WHO showed no or little effect in hospitalized COVID-19 patients, including metrics of overall mortality, initiation of ventilation and duration of hospital stay ^18^. Thus, therapeutic efficacy of remdesivir appears far from satisfactory, and new therapeutic options are urgently needed.

The current work started through development of a target-specific cell-based screening assay. We created the Mpro sensor, and our screen of 1622 FDA-approved drugs led to identification of 9 drugs that inhibit Mpro activity and show anti-SARS-CoV-2 activity. We characterized the antiviral property of five drugs further. Four of them, including simeprevir, ABT-199, hydroxyprogesterone and cinacalcet effectively inhibited Mpro and blocked SARS-CoV-2. This suggests that these latter four drugs inhibit SARS-CoV-2 mainly via inhibition of Mpro activity. These drugs have similar or better EC50 than ebselen, and thus they may be repurposed as potential therapies against COVID-19. Furthermore, previous computational docking predicted that simeprevir and venetoclax (ABT-199) bind and inhibit Mpro ^19,20^. Hydroxyprogesterone inhibits SARS-CoV-2, but its mode of action is unclear ^21^. Our data suggests that hydroxyprogesterone may block SARS-CoV-2 by inhibiting Mpro activity. Utilizing our FlipGFP-based HTS approach enables screening of Mpro inhibitors in the cellular context, and can be expanded to visualize activity of key proteases of other viruses at BSL2, and to identify their inhibitors.

The most potent antiviral drug discovered in the current work, ethacridine, showed higher antiviral potency than remdesivir and very little cell toxicity (**Fig. 3e**). This agent blocks SARS-CoV-2 mainly by inactivating the viral particles. First, unlike the other identified drugs that show similar range of IC_50_ in inhibiting Mpro and EC50 against SARS-CoV-2, ethacridine has much higher anti-viral potency (EC_50_ ∼ 0.08 *μ*M) than its Mpro-inhibiting activity (IC_50_ ∼ 3.5 *μ*M), indicating that the main mode of action of ethacridine is not via inhibition of Mpro. Second, we provide direct evidence that after ethacridine treatment, SARS-CoV-2 had markedly reduced infectivity to bind to host cells. On the other hand, ethacridine does not block viral RNA replication as shown by the qRT-PCR data. Therefore, our data suggest that ethacridine blocks SARS-CoV-2 mainly by inactivating the viral particles without influencing viral RNA replication. The precise mechanisms for how ethacridine inactivates the viral particles infection will require further investigation, such as potential ultrastructural changes of viral particles and/or binding of ethacridine to viral RNA and/or protein.

Our findings herein reveal a new approach against SARS-CoV-2 through the inactivation of viral particles, for which the efficacy is expected to be cell type-independent. Indeed, ethacridine blocks SARS-CoV-2 infection of human cell line A549^ACE2^ cells and also in primary HNE cells. Furthermore, ethacridine is non-toxic in various animal models including rat, mice and rabbits. For example, mice treated with 20 mg/kg ethacridine by i.p. injection showed no toxicity ^22^. While currently ethacridine is mainly used as a topical wound disinfectant, it has been applied to patients for treating puerperal sepsis via intravenous injection ^23^. Thus, it is plausible to validate its antiviral effect in animal models and COVID-19 patients. Moreover, because it blocks the coronavirus by direct inactivation of the viral particles, ethacridine may be used in combination with other pathway-targeted drugs such as the replicase inhibitor remdesivir in treating COVID-19 patients for potentially greater clinical efficacy.

## Methods

### Plasmid construction

Plasmid constructs were created by standard molecular biology techniques and confirmed by exhaustively sequencing the cloned fragments. To create pcDNA3-Mpro-flipGFP-T2A-mCherry, the GFP10-E5-GFP11-Mprosubstrate-K5 fragment was first generated by PCR with an overlapping primer with coding sequence of the MPro substrate, AVLQSGFR. This fragment was then digested with NheI and AflII and inserted into the pcDNA3-TEV-flipGFP-T2A-mCherry ^11^.

### Characterization of GFP-based Mpro activity sensor FlipGFP^Mpro^

To check the FlipGFP^Mpro^ signal in the presence of wildtype or mutant MPro of SARS-CoV-2, HEK293T cells were seeded onto eight-well plate and 24 hours later, 40 ng of pcDNA3-FlipGFP^Mpro^-T2A-mCherry was transfected with 40 ng of pLVX-EF1alpha-nCoV2019-nsp5-2xStrep-IRES-Puro (expressing wildtype Mpro of SARS-CoV-2) or pLVX-EF1alpha-nCoV2019-nsp5-C145A-2xStrep-IRES-Puro (expressing C145A MPro mutant). To check the FlipGFP^Mpro^ signal after SARS-CoV-2 infection, HEK293T on 10-mm coverglasses were transfected with 200 ng of pcDNA3.1-hACE2 (Addgene, #145033) 24 hours later. Those 293T cells were further infected with SARS-CoV-2 at an MOI of 0.5. At 24 hours after infection, 293T cells were fixed and processed for immunostaining against dsRNA (Scicons, 10010500) and visualized with goat anti-mouse IgG H&L (Cy5 ®) (Abcam, ab6563). Images were taken using Nikon Eclipse Ti-E Spinning Disk under 20X and 60X. The cell pixel intensity in the GFP and mCherry channels were scored using Analyze Particle function in ImageJ. The ratio of GFP pixel intensity against mCherry pixel intensity was then compared in cells with wildtype and mutant MPro or in cells with SARS-CoV-2 or mock infection.

### High-throughput screen (HTS) against the FDA-approved library

An FDA-approved drug library (MedChemExpress) containing 1622 compounds was used for HTS. For initial screening, 293T cells were seeded onto a 96-well plate, incubated overnight and transfected with 20 ng of pcDNA3-Mpro-flipGFP-T2A-mCherry and 20ng pLVX-EF1alpha-nCoV2019-nsp5-2xStrep-IRES-Puro encoding MPro using calcium-phosphate transfection. At 3 hours after transfection, compounds were added to individual wells at a 100X dilution for a final concentration of 20 *μ*M. Verification of the 120 hits from the initial screening and further IC_50_ testing of the 12 hits followed a similar protocol except that a decreasing concentration series was used. Images were acquired 20 hours after transfection in the GFP and mCherry channels with a Nikon Eclipse Ti-E Spinning Disk. The cells’ pixel intensity in the GFP and mCherry channels were scored using Analyze Particle function in imageJ. MPro activity was calculated as the ratio of the GFP pixel intensity versus the mCherry pixel intensity.

### Antiviral assay

The African green monkey kidney Vero E6 cell line was obtained from the American Type Culture Collection (ATCC, no. 1586) and maintained in Minimum Essential Medium (MEM; Gibco Invitrogen) supplemented with 10% fetal bovine serum (FBS; Gibco Invitrogen), 1% penicillin-streptomycin-glutamine (Gibco Invitrogen), at 37°C in a humidified 5% CO_2_ incubator. A clinical isolate of SARS-CoV-2 (USA-WA1/2020, BEI Cat No: NR-52281) was propagated in Vero E6 cells. Viral titers were quantified with a plaque assay. All the infections were performed at biosafety level-3 (BSL-3).

To assess the antiviral activity, ∼70% confluent monolayers of Vero E6 cells (3×10^5^ cells/well in 24-well plates) were pretreated with drugs at different concentration for 3 hours (pretreatment) and then infected with SARS-CoV-2 (MOI=0.5) at 37°C for 1 hour. The virus solution was removed, cells were further cultured with fresh medium containing drugs at different concentrations. At 16 hours post-infection, viral titers of the supernatants were detected with a plaque assay.

### Plaque assay

Confluent monolayers of Vero E6 cells grown in six-well plates were incubated with the serial dilutions of virus samples (250 μl/well) at 37°C for 1 hour. Next, the cells were overlayed with 1% agarose (Invitrogen) prepared with MEM supplemented with 2% FBS. Three days later, cells were fixed with 4% formaldehyde for 2 hours, the overlay was discarded, and samples were stained with crystal violet dye.

### Viral RNA quantification

Viral RNA (vRNA) was extracted from cell pellets or supernatants with Tri-reagent (Ambion), following the manufacturer’s instructions. The supernatant samples were pretreated with RNAse A for 2 hours at 37°C to remove non-encapsidated RNA. These samples were also spiked with equivalent volumes of *Drosophila* C Virus (DCV) to enable normalization, based on variation in RNA extractions. RNA from supernatants was used directly to make cDNA using the iScript RT Supermix (BioRad). For cell pellets, 1–2 mg of RNA was treated with DNAse I (NEB), x µL and 2 µL of this reaction was used to make cDNA. qPCR was done using the Luna Universal qPCR Master Mix (NEB) and run on a CFX connect qPCR Detection System (BioRad). To determine the number of vRNA copies per ml, plasmids containing the nucleocapsid gene of SARS-CoV-2 (cloned from the USA-WA1/2020 isolate) or the DCV full genome were used as standards and diluted serially 10-fold to determine target copy numbers. Threshold cycle (Ct) values were plotted against the number of target-copies, and the resultant standard curve was used to determine the number of genome equivalents of vRNA in the samples. For cell pellet samples, the vRNA copy number was normalized to the housekeeping gene *huel*, and supernatant samples were normalized to spiked DCV RNA. All samples were within the range of linearity of a standard curve and primers efficiencies were 100% +/- 5%. The primers used for SARS-CoV-2 are 5′-TCCTGGTGATTCTTCTTCAGG-3′ and 5′-TCTGAGAGAGGGTCAAGTGC-3′, DCV 5-CAGCAAAGAAACAGCGTGAG-3’ and 5′-CACTTGCGCAACAATACGAG-3′, huel 5′-TCAGACGACGAAGTCCCCATGAAG-3′ and 5′-TCCTTACGCAATTTTTTCTCTCTGGC-3′

### Cytotoxicity assay

Cytotoxicity of the identified drugs on Vero E6 cell was determined with WST-1 cell proliferation assays (ROCHE, 5015944001). Twenty thousand cells were seeded into a 96-well plate and incubated for 20–24 h at 37 °C, and 1 *μ*L of each compound at decreasing concentrations was added. After 18 hrs incubation at 37 °C, WST-1 assays were performed according to manufacturer’s protocols. All experiments were performed in triplicate.

### Adsorption assay

The virus sample was supplemented with 5 *μ*M ethacridine or with DMSO and incubated for 1 hour at 37°C. Samples were added to ∼70% Vero E6 cells monolayers on coverslips with an MOI of 100. The cells were immediately placed on ice for 1 hour, and then the virus suspension was quickly removed, and cells were washed three times with ice-cold PBS. Cells were immediately fixed with 4% PFA for 30 min at room temperature. PFA was further washed with PBS and quenched with 1 M glycine in PBS. Immunostaining was further carried out using an antibody against nucleocapsid (Genetex, SARS-CoV-2 (COVID-19) nucleocapsid antibody, GTX135357) or an antibody against spike (Genetex, SARS-CoV-2 (COVID-19) spike antibody [1A9], GTX632604) and visualized with goat anti-rabbit IgG H&L (Alexa Fluor® 488) (Abcam, ab150077) or goat anti-mouse IgG H&L (Alexa Fluor® 555) (Abcam, ab150114). Images were acquired using Nikon Eclipse Ti-E Spinning Disk under 60X and processed in ImageJ. Fluorescence signals were calculated by pixel intensity using Analyze Particle function in Image J.

### Statistics

All statistics were performed in GraphPad Prism. IC_50_ or EC_50_ of MPro inhibition, cytotoxicity and antiviral activity was calculated using the non-linear fit function (Variable slope). Non-paired t-test was used to compare differences between groups. One way-ANOVA and Dunnett’s multiple comparisons test was used to compare differences among multiple groups.

## Acknowledgments

We thank David Gordon and Nevan Krogan for sharing the cDNA of Mpro of SARS-CoV-2.

## Author contributions

X.L., X.S. initiated the project and designed the fluorescent assays. X.L., J.Y. prepared the drug library. X.L. performed screening and characterized dose response of Mpro inhibition. P.L., Y.X., R.A. designed the antiviral activity assays. P.L., Y.X. conducted plaque assay. P.L., Y.X., X.L. performed quantitative analysis. M.K. performed RT-qPCR. X. L. performed toxicity assays. P.L., Y.X., X.L. performed and analyzed immuno-staining. P.L., Y.X., T.N., J.V.N., C.W. P. K. J. and R.A. designed the HNE cell infection experiments. P.L. and Y.X. performed viral infection, C.W., T.N. performed cell culturing, C.W. conducted immunostaining and imaging, X.L. performed quantitative analysis of the HNE cells. X.S. wrote initial manuscript, X.L., P.L., Y.X., M.K., C.W. wrote the methods. All authors edited and contributed to the final draft.

## Data and materials availability

All data are available in the main text or in the supplementary information.

## Supplementary Information

Supporting Figs. S1–6

---

**Supporting Fig. S1.**
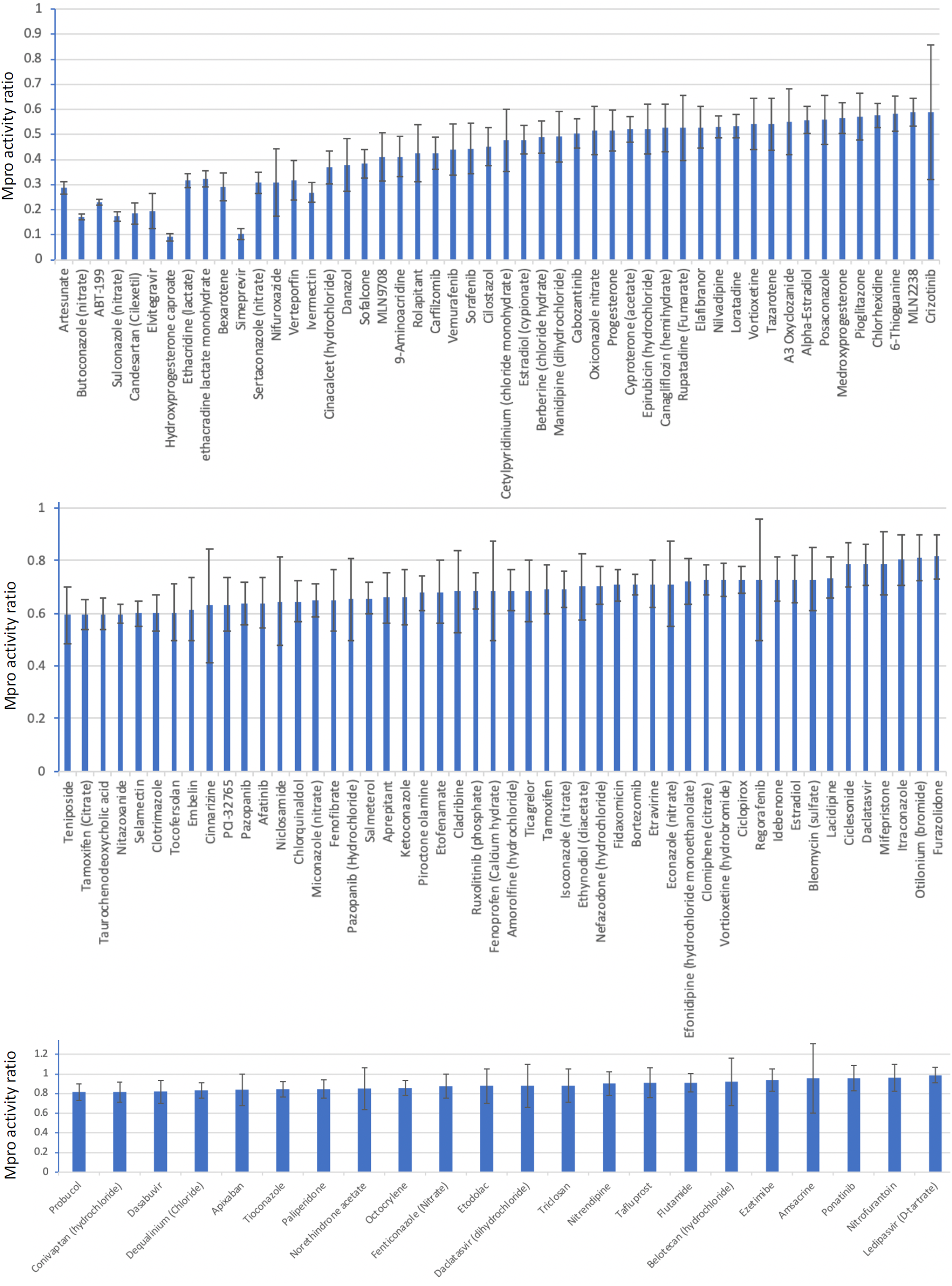
Verification of the selected top 120 drugs using the Mpro activity reporter FlipGFP^Mpro^. Ratio of Mpro activity was calculated based on FlipGFP^Mpro^ fluorescence of drug-incubated HEK293 cells, divided by that of DMSO-treated HEK293 cells. FlipGFP^Mpro^ fluorescence was normalized by mCherry in HEK293 cells, which co-expressed FlipGFP^Mpro^, mCherry, and Mpro. Data are mean ± SD (n = 5).

**Supporting Fig. S2.**
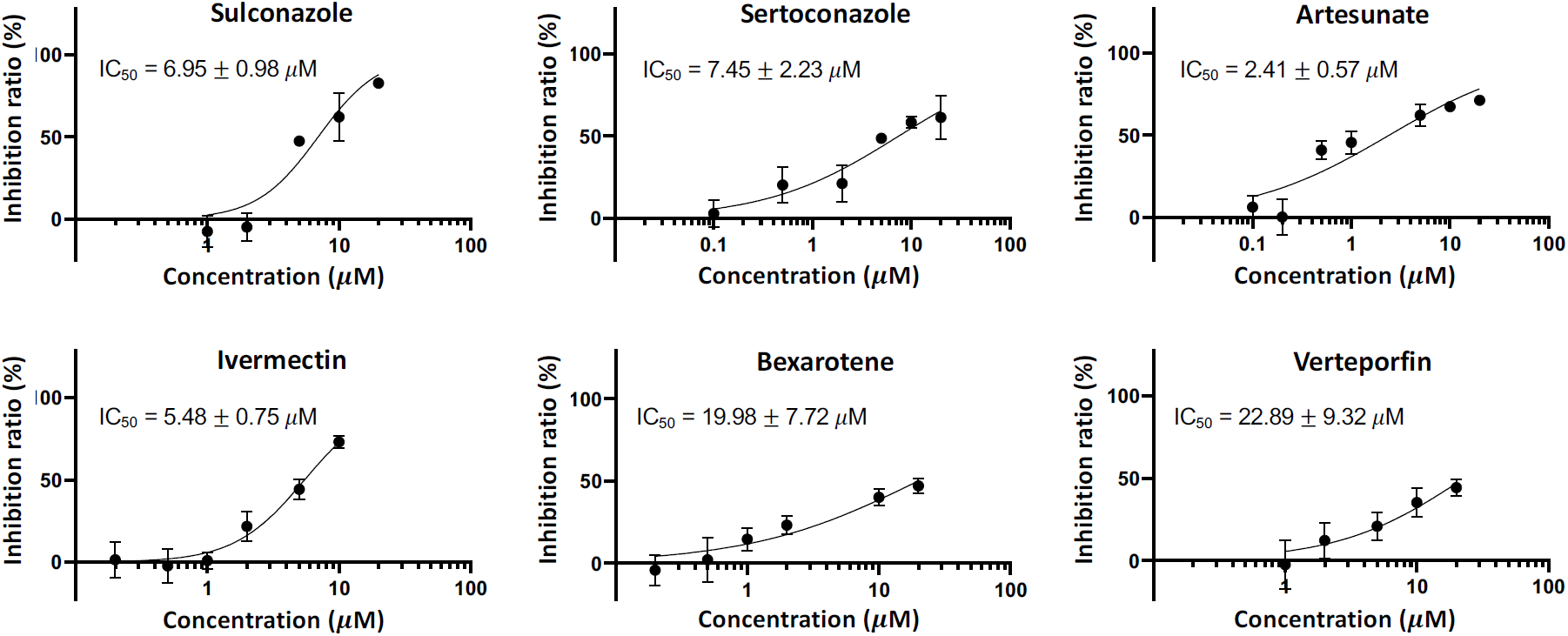
Dose-response curve of Mpro inhibition. The Mpro activity was determined as FlipGFP fluorescence normalized by mCherry. The ratios of Mpro activity were calculated by normalizing Mpro activity with that of cells treated with DMSO. Inhibition ratio was calculated as (1-(ratio of Mpro activity)) X100%. Data are mean ± SD (n = 5). IC_50_ was represented as mean ± SEM (n = 5).

**Supporting Fig. S3.**
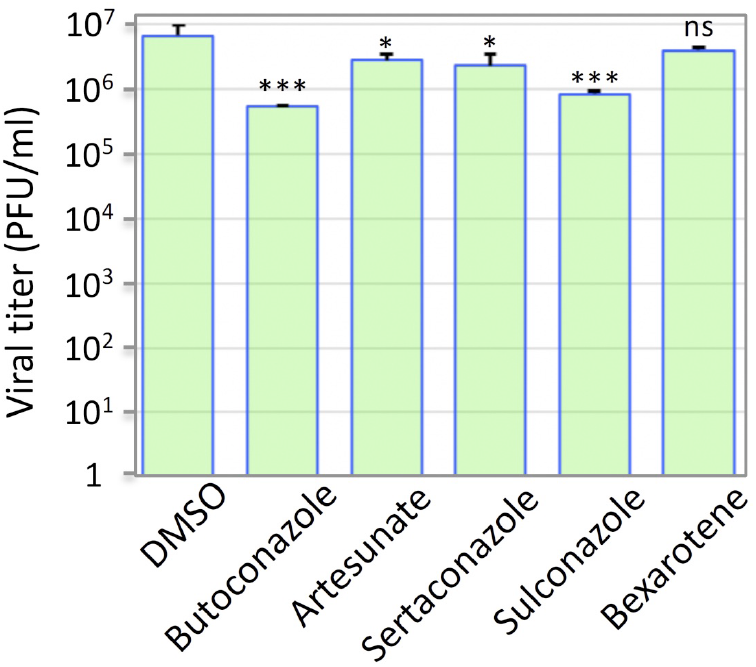
Antiviral activities of the identified drugs. Antiviral activities of five drugs (5 *μ*M) were quantified by a plaque assay with SARS-CoV-2 in Vero E6 cells. Data are mean ± SD (n = 3). *: p value < 0.05; ***: p value < 0.001. ns: not significant. PFU: plaque-forming unit.

**Supporting Fig. S4.**
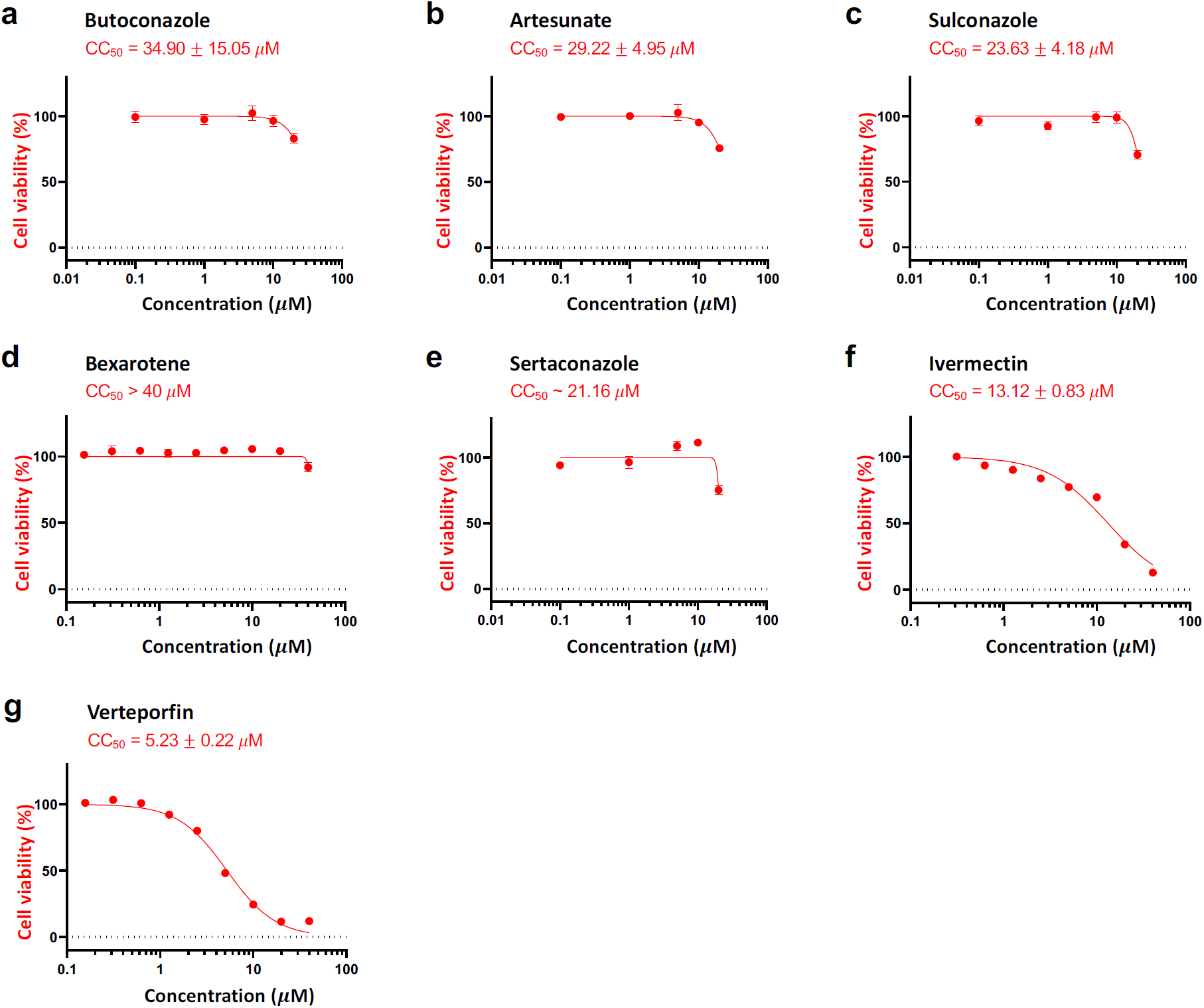
Cytotoxicity of the identified drugs. The cytotoxicities of the indicated compounds were determined in Vero E6 cells with the WST-1 assay. CC_50_ is represented as mean ± SEM (n = 3).

**Supporting Fig. S5.**
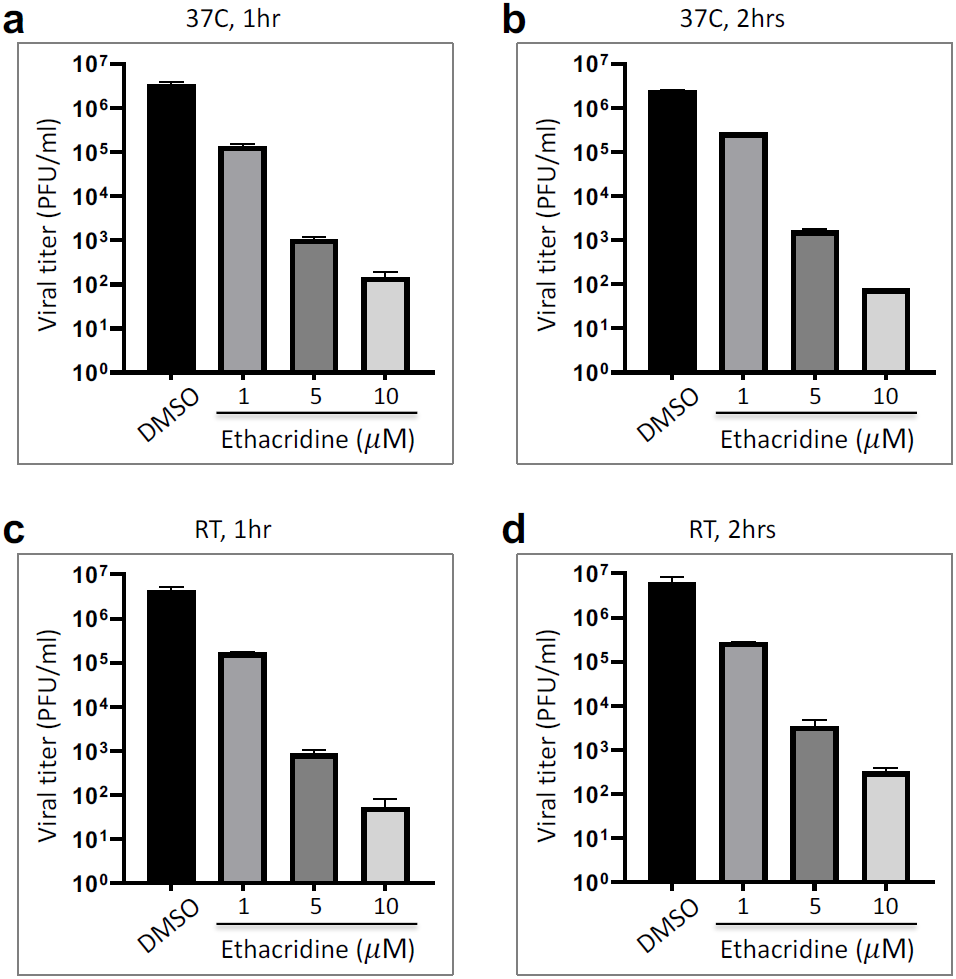
Virucide effect of ethacridine on SARS-CoV-2. Effects of ethacridine on the infectivity of SARS-CoV-2 were examined using plaque assay at 37°C (**a, b**) or in the room temperature (RT) (**c, d**). SARS-CoV-2 was mixed with ethacridine for 1 or 2 hours before being added to infect Vero E6 cells. Data are mean ± SEM (n = 3).

**Supporting Fig. S6.**
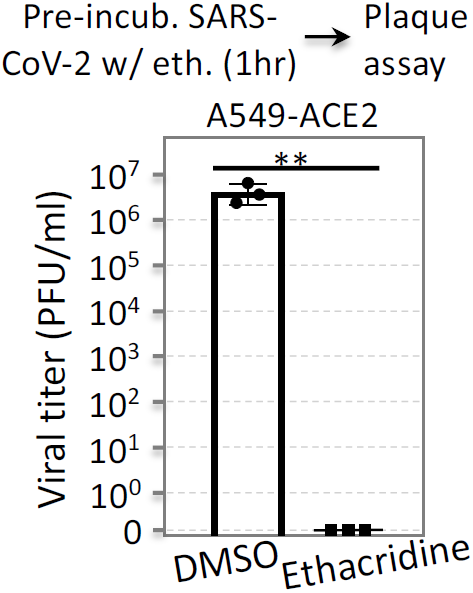
Quantitative analysis of viral titer by plaque assay in the human cells A549 that stably express ACE2. SARS-CoV-2 was pre-incubated with ethacridine (5 uM) for 1hr, followed by plaque assay on the human A549 cells stably expressing human ACE2 (A549^ACE2^).

